# Mutant RIT1 cooperates with YAP to drive an EMT-like lung cancer state

**DOI:** 10.1101/2024.11.11.623044

**Authors:** Mary C. Rominger, Saksham Gupta, Sitapriya Moorthi, Maria McSharry, Shriya Kamlapurkar, Siobhan O’Brien, Annie Waldum, April Lo, Fujiko Duke, Amy R. Lowe, Elizabeth Cromwell, Raisa Glabman, Amanda Koehne, Alice H. Berger

## Abstract

The discovery of oncogene addiction in cancer has led to the development of over a dozen FDA-approved biomarker-driven therapies in lung adenocarcinoma. Somatic mutations of the “Ras-like in all tissues” (RIT1) gene are non-canonical driver events in lung cancer, occurring in ∼2% of lung adenocarcinomas in a mutually exclusive fashion with *KRAS* and *EGFR* mutations. Patients with *RIT1*-mutant lung cancer lack targeted therapy treatment options, and a lack of pre-clinical models has hindered the development of therapeutic strategies for *RIT1*-mutant lung cancer. Here we report a new mouse model of RIT1-driven lung cancer in which the human RIT1^M90I^ variant can be induced in a Cre-regulated manner. We show that autochthonous expression of RIT1^M90I^ in the lung weakly promotes cancer alone or in combination with loss of the *p53* tumor suppressor. However, potent synergy between RIT1^M90I^ and inactivation of *Nf2* drives an aggressive epithelial-to-mesenchymal (EMT) lung cancer with 100% penetrance and short latency. We show this oncogenic cooperation is driven by synergistic activation of cJUN, a component of the AP-1 complex. Therapeutic inhibition of MEK and YAP/TEAD suppressed RIT1-driven lung cancer in vivo. These data identify YAP/TEAD as an important mediator of RIT1’s oncogenic potential and nominate TEAD as an important drug target in *RIT1*-mutant lung cancer.

**HIGHLIGHTS:**

- We report a new RIT1^M90I^-mutant autochthonous lung tumor model
- The most common oncogenic variant of RIT1, RIT1^M90I^, weakly promotes lung tumor development
- RIT1^M90I^ drives the formation of lethal lung tumors in cooperation with *p53* and *Nf2* tumor suppressor gene loss
- RIT1^M90I^ and YAP cooperatively regulate cJUN expression
- Therapeutic MEK and TEAD targeting suppresses RIT1^M90I^-driven tumorigenesis

## INTRODUCTION

Aberrant activation of genes in the EGFR/Ras pathway appears to be central to the development and progression of non-small cell lung cancer and its most common histological subtype, lung adenocarcinoma (TCGA 2014). Following the discovery of somatic *EGFR* mutations in lung adenocarcinoma in 2004 (Paez et al. 2004; Pao et al. 2004; Lynch et al. 2004), it was recognized that over ten other genes related to EGFR signaling are also commonly mutated such as Ras genes, *BRAF, ERBB2, MET*, and fusions of receptor tyrosine kinase (RTK) genes (Imielinski et al. 2012; TCGA 2014; Campbell et al. 2016; Harada et al. 2023). This has spurred remarkable cancer therapeutic development resulting in FDA approval of dozens of targeted inhibitors of proteins in the EGFR signaling pathway (Melosky et al. 2021), including the 3rd generation covalent inhibitor of EGFR, osimertinib (Wu et al. 2020; Soria et al. 2017), and cysteine-selective covalent KRAS G12C inhibitors sotorasib and adagrasib (Hong et al. 2020; Jänne et al. 2022). However, many lung adenocarcinomas do not harbor mutations in these druggable genes, limiting targeted therapy options for the remaining one third or >30,000 lung adenocarcinoma patients in the U.S. every year.

*RIT1* is a Ras-family GTPase gene with significant homology to *HRAS, NRAS*, and *KRAS*. Like these canonical Ras genes, *RIT1* is a recurrent target of somatic mutations and copy number amplification in cancer, particularly lung cancer *(Van et al. 2020; Berger et al. 2014)*. We previously identified somatic mutations of *RIT1* in 2% of lung adenocarcinomas (TCGA 2014; Berger et al. 2014; Imielinski et al. 2012). An additional 24% of lung adenocarcinomas have amplification or overexpression of *RIT1* (TCGA 2014) (**Figure S1A-B**). Also reminiscent of Ras proteins, germline mutations in *RIT1* cause the “RASopathy” Noonan syndrome (Aoki et al. 2013). In both cancer and Noonan syndrome, RIT1 variants are enriched near the Switch II pocket of the GTPase, but unlike Ras proteins, RIT1 variants appear to function via induction of increased RIT1 protein abundance (Castel et al. 2019; Lo et al. 2021). Germline mutation of *Rit1* in the mouse phenocopies human Noonan syndrome (Castel et al. 2019), and somatic activation of Rit1^M90I^ in the hematopoietic compartment results in myeloproliferative disorder (Chen et al. 2022). However, the lack of murine and patient-derived models of mutant *RIT1* function in lung cancer has limited discovery of therapeutic approaches in *RIT1*-mutant lung cancer. Because *RIT1* mutations are mutually exclusive with other EGFR-pathway driver oncogenes, there is an unmet medical need to develop strategies to target *RIT1*-mutant tumors. Availability of a mouse model of *RIT1*-mutant lung cancer would enable new advances for this subset of oncogene-driven lung cancers.

Here we describe the generation of an autochthonous genetically-engineered mouse model of RIT1^M90I^-mutant lung cancer in which Cre expression induces excision of a lox-STOP-lox cassette and transgenic expression of RIT1^M90I^, the most recurrent cancer-associated variant of RIT1 (**Figure 1A** and **Figure S1C**). RIT1^M90I^ induction in the murine lung induces weakly penetrant lung tumor formation at long latency. However, we find that combined inactivation of *p53* and *Nf2* tumor suppressors synergizes with RIT1^M90I^ expression to drive an aggressive epithelial-to-mesenchymal transition (EMT) lung cancer state. Activation of YAP/TEAD, either by engineered YAP1 mutation or by *Nf2* loss, synergizes with RIT1^M90I^ to drive an oncogenic program involving the AP-1 transcription factor cJUN and resulting tumor cell growth is both MEK- and TEAD-dependent. These data identify MEK and TEAD as important mediators of RIT1’s oncogenic function and present a path forward for targeted therapy development in *RIT1*-mutant lung cancer.

**Figure 1.**
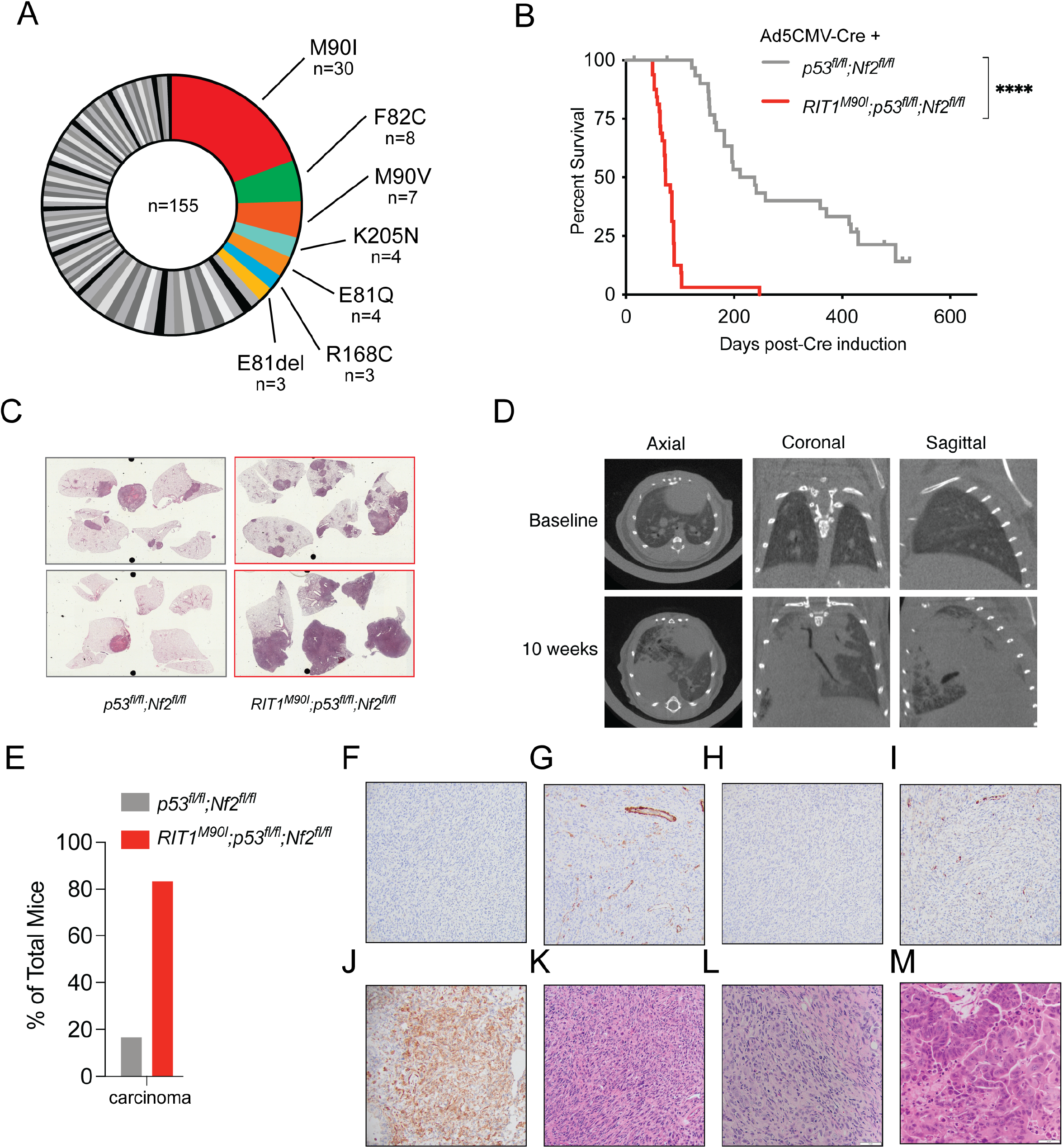
RIT1^M90I^ synergizes with *p53* and *Nf2* loss to drive lung tumorigenesis. **A**, Mutations in *RIT1* from non-small cell lung cancer tumors in AACR GENIE v16.0. **B**, Kaplan-Meier curve of overall survival of *RIT1*^*M90I*^*/p53*^*fl/fl*^*/Nf2*^*fl/fl*^ (n=32) and *p53*^*fl/fl*^*/Nf2*^*fl/fl*^ animals (n=32) following intratracheal Adeno-CMV-Cre delivery. **C**, Representative H&E staining of lungs from *RIT1*^*M90I*^*/p53*^*fl/fl*^*/Nf2*^*fl/fl*^ and *p53*^*fl/fl*^*/Nf2*^*fl/fl*^ mice at the humane endpoint. **D**, Representative microCT images of lungs from a *RIT1*^*M90I*^*/p53*^*fl/fl*^*/Nf2*^*fl/fl*^ mouse prior to and 10 weeks post-Ad5CMV-Cre delivery. **E**, Pathologist-determined carcinoma incidence in *RIT1*^*M90I*^*/p53*^*fl/fl*^*/Nf2*^*fl/fl*^ (n=6) and *p53*^*fl/fl*^*/Nf2*^*fl/fl*^ animals (n=6) at 6-9 weeks post Ad5CMV-Cre delivery. **F-J**, Immunohistochemistry staining of a *RIT1*^*M90I*^*/p53*^*fl/fl*^*/Nf2*^*fl/fl*^ showing negative staining of antibodies against mesothelin (F), smooth muscle actin (G), cytokeratin (H), and desmin (I), and positive staining for vimentin (J). **K-M**, H&E staining showing regions of sarcomatoid (K), undifferentiated (L), and adenocarcinoma (M) morphology.

## RESULTS

### RIT1^M90I^ synergizes with *p53* and *Nf2* loss to drive lung tumorigenesis

To enable in vivo study of mutant RIT1 function, we generated gene-targeted mice harboring a lox-stop-lox-RIT1^M90I^ cassette in the *Rosa26* locus under endogenous control of the Rosa26 promoter (Methods; **Figure S1D-E**). B6 founder mice were intercrossed to 129Sv/ImJ wild-type mice and the line was maintained on a mixed genetic background. Germline activation of RIT1^M90I^ resulted in embryonic lethality, possibly due to differences in Rosa26 expression regulation compared to normal endogenous regulation of murine Rit1 (**Figure S1F**). To generate an autochthonous model of lung-specific RIT1^M90I^ activation, we delivered Cre recombinase to the lung of heterozygous LSL-RIT1^M90I^ mice and wild-type littermate controls using intratracheal Ad5CMV-Cre delivery (DuPage et al. 2009). While recombination of the targeted allele and expression of the human transgene was readily observed in the Cre-treated lungs of LSL-RIT1^M90I^ mice (**Figure S1G-H**), there was no significant difference in overall survival (**Figure S1I**) of RIT1^M90I^ mice compared to littermate controls. A modestly increased adenoma prevalence was observed in the RIT1^M90I^-positive mice compared to wild-type controls (**Figure S1J**).

In human lung adenocarcinoma, *TP53* mutations co-occur with *RIT1* mutations in over 50% of *RIT1*-mutant tumors (**Figure S2A**), and deletion of murine *Trp53* (hereafter *p53*) is often a requisite cooperating event in murine models of lung cancer. To determine whether RIT1^M90I^ could drive tumor formation with combined *p53* deletion, we generated cohorts of LSL-RIT1^M90I^ heterozygotes in the presence and absence of a conditional Cre-regulated *p53* allele. Surprisingly, no significant difference in either survival (**Figure S2B**) or adenoma/carcinoma incidence (**Figure S2C**) was observed between LSL-RIT1^M90I^ animals with intact *p53* (*p53*^*WT*^) or homozygous loss of p53 (*p53*^*fl/fl*^). These data contrast with the rapid development of lung adenocarcinomas in murine models of *Kras*-mutant lung cancer (Jackson et al. 2001) and suggest that RIT1^M90I^ may require additional cooperating events to drive tumorigenesis.

We previously performed genome-wide CRISPR dependency mapping to identify gene knockouts that either suppress RIT1^M90I^-driven phenotypes or synergize with RIT1^M90I^ to promote oncogenic phenotypes (Vichas et al. 2021). We showed that loss of Hippo-pathway genes potently synergized with RIT1 in models of drug resistance and cancer growth (Vichas et al. 2021). Importantly, human *RIT1*-mutant lung adenocarcinomas also show suppression of Hippo pathway gene expression and increased nuclear YAP accumulation, suggesting that Hippo inactivation and YAP activation may be important co-occurring factors in human RIT1-mutant lung cancer (Vichas et al. 2021).

To model the combined Hippo loss and *RIT1* mutation observed in human lung adenocarcinoma, we tested whether Hippo pathway inactivation via *Nf2* deletion would modify the ability of RIT1^M90I^ to promote lung cancer. Remarkably, in the context of combined *Nf2/p53* deletion, RIT1^M90I^ drove an aggressive lung cancer phenotype, with >95% of animals succumbing to disease within 100 days and 100% penetrance at <1 year (**Figure 1B-C**) with significantly shorter latency than tumor formation in *p53*^*fl/fl*^*/Nf2*^*fl/fl*^ controls (**Figure 1B**; p < 0.0001 by log-rank test). Tumor progression could be monitored by microCT imaging (**Figure 1D** and **Figure S2D**). At only 6-9 weeks post-Cre delivery, 83% of *RIT1*^M90I^/*p53*^*fl/fl*^*/Nf2*^*fl/fl*^ animals had lung carcinomas, compared to 16.3% of *p53*^*fl/fl*^*/Nf2*^*fl/fl*^ (**Figure 1E**). Loss of *p53* appeared to be required for tumor formation, as survival was significantly extended in the presence of intact p53 (**Figure S2E**). Tumors were negative by immunohistochemistry for mesothelin, smooth muscle actin, pan-cytokeratin, and desmin, and positive for vimentin (**Figure 1F-J**). Tumors appeared undifferentiated with areas of sarcomatoid or adenocarcinoma histology also apparent in some cases (**Figure 1K-M**). These data demonstrate that RIT1^M90I^ cooperates with loss *p53/Nf2* to promote lung tumorigenesis in vivo.

### Expression of RIT1^M90I^ in alveolar epithelial cells drives an EMT-high lung cancer state

In lung cancer, *RIT1* mutations are found primarily in lung adenocarcinoma. The tumors observed in the *RIT1*^M90I^/*p53*^*fl/fl*^*/Nf2*^*fl/fl*^ lung model, however, appeared more mesenchymal-like, resembling sarcomas or mesotheliomas but negative for markers of those tumor types such as mesothelin (**Figure 1F**). The cell-of-origin of lung adenocarcinoma is thought to be the alveolar type II (AT2) cell (Ferone et al. 2020). Because the Ad5CMV-Cre virus used to initiate tumor formation can infect any cell type of the lung, we hypothesized that restricting RIT1^M90I^ to the AT2 compartment might drive a tumor phenotype with greater resemblance to human lung adenocarcinoma. We therefore next generated cohorts of *RIT1*^M90I^/*p53*^*fl/fl*^*/Nf2*^*fl/fl*^ and *p53*^*fl/fl*^*/Nf2*^*fl/fl*^ mice and delivered Cre via intratracheal delivery of Ad5mSPC-Cre, which provides AT2-specific Cre expression under control of the surfactant protein C (SPC) promoter (Sutherland et al. 2011). Surprisingly, similar to mice treated with Ad5CMV-Cre, delivery of Ad5mSPC-Cre to *RIT1*^M90I^/*p53*^*fl/fl*^*/Nf2*^*fl/fl*^ mice resulted in rapid development of lung tumors (**Figure 2A**) with similar histopathology to the mice treated with Ad5mCMV-Cre (**Figure S3A-C**). We derived cell lines from one *p53*^*fl/fl*^*/Nf2*^*fl/fl*^ tumor, two Ad5mSPC-Cre/*RIT1*^M90I^/*p53*^*fl/fl*^*/Nf2*^*fl/fl*^ tumors, and one Ad5CMV-Cre/*RIT1*^M90I^/*p53*^*fl/fl*^*/Nf2*^*fl/fl*^ tumor (**Figure 2B**). Western blot confirmed robust protein expression of RIT1 in the three *RIT1*^*M90I*^ tumors compared to the *p53*^*fl/fl*^*/Nf2*^*fl/fl*^ control tumor cells (**Figure 2B**).

**Figure 2.**
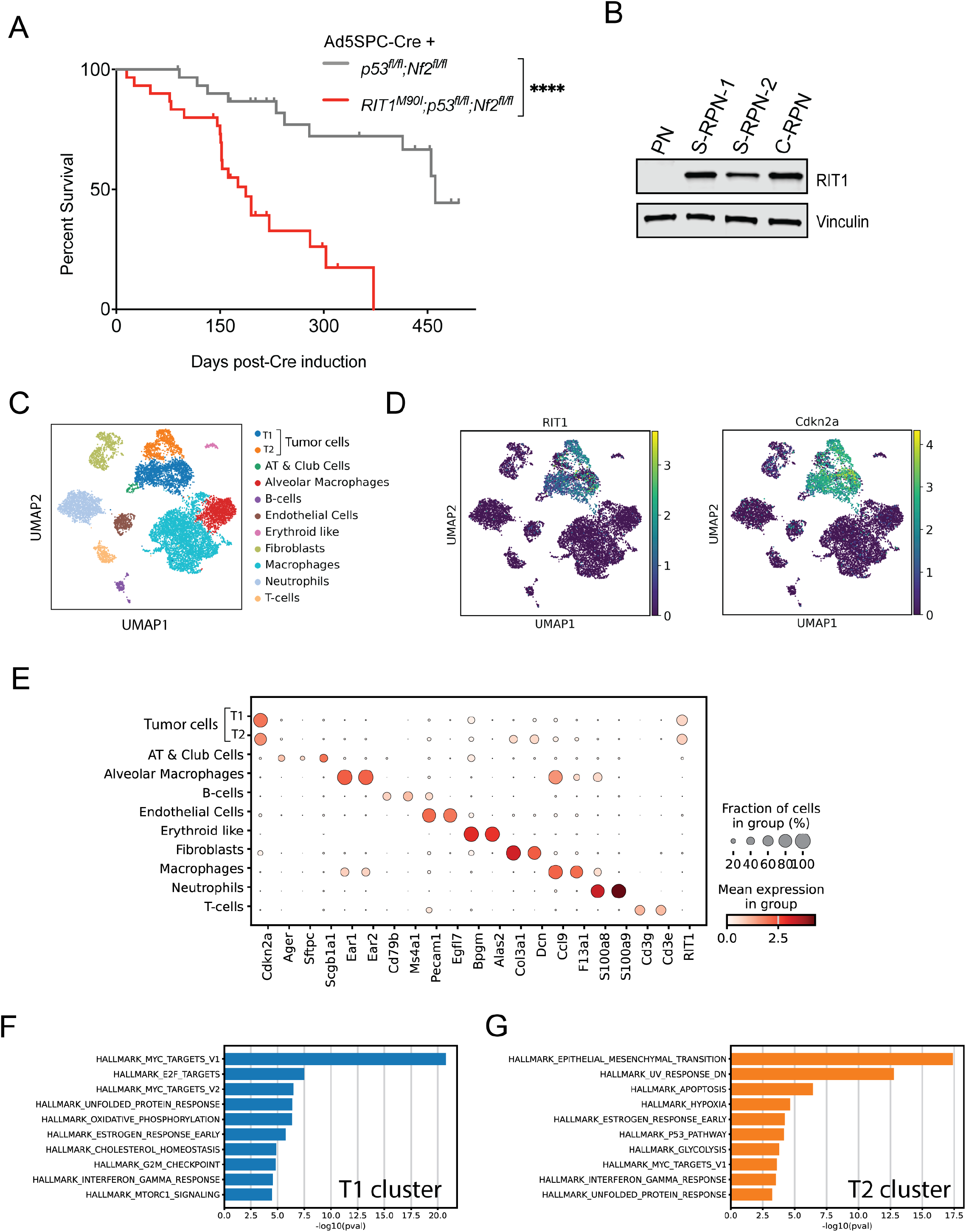
Expression of RIT1^M90I^ in alveolar epithelial cells drives an EMT-high lung cancer state. **A**, Kaplan-Meier curve of overall survival of *RIT1*^*M90I*^*/p53*^*fl/fl*^*/Nf2*^*fl/fl*^ (n=30) and *p53*^*fl/fl*^*/Nf2*^*fl/fl*^ animals (n=31) following intratracheal Ad5mSPC-Cre delivery. **B**, Western blot of RIT1 and Vinculin (loading control) in cell lines derived from tumors from Ad5CMV-Cre/*p53*^*fl/fl*^*/Nf2*^*fl/fl*^ (“PN”), Ad5mSPC-Cre/*RIT1*^*M90I*^*/p53*^*fl/fl*^*/Nf2*^*fl/fl*^ *(“S-RPN-1” and “S-RPN-2*”), or Ad5CMV-Cre/*RIT1*^*M90I*^*/p53*^*fl/fl*^*/Nf2*^*fl/fl*^ mice. **C**, UMAP clustering of scRNA-seq data of cells derived from a Ad5mSPC-Cre/*RIT1*^*M90I*^*/p53*^*fl/fl*^*/Nf2*^*fl/fl*^ tumor 20 weeks after viral delivery. **D**, UMAP clustering of the same data from (C), with relative gene expression of the human RIT1^M90I^ transgene (left) or *Cdkn2a* (right) indicated. **E**, Dot plot showing expression of marker genes used for cell type assignments (Methods). **F**, GSEA of the top 50 differentially expressed genes from tumor cluster 1 (T1). See also **Table S1. G**, GSEA of the top 50 differentially expressed genes from tumor cluster 2 (T2). See also **Table S1**.

To better understand the lineage phenotype of *RIT1*^*M90I*^-mutant tumors, we performed single-cell RNA-sequencing on cells isolated from a tumor from a moribund Ad5mSPC-Cre-treated *RIT1*^M90I^/*p53*^*fl/fl*^*/Nf2*^*fl/fl*^ animal (**Figure 2C**). Clustering of tumor and stromal cells identified 11 different cell clusters, 9 of which could readily be linked to known cell types such as T cells, alveolar macrophages, and alveolar epithelial cells (**Figure 2C**). Two clusters could not be matched to known cell types, and alignment of reads to human mutant *RIT1*^*M90I*^ revealed that transgene expression was restricted to these two related clusters, identifying these cells as the tumor cells (**Figure 2D-E; Table S1**). In addition, both tumor clusters demonstrated robust upregulation of the tumor suppressor *Cdnk2a* (**Figure 2D-E**), likely a consequence of feedback regulation after p53 loss (Sherr 2001; Stott et al. 1998). Gene set enrichment analysis (GSEA) of Tumor cluster 1 (T1) showed elevated proliferative markers such as MYC and E2F targets as well as keratins typical of lung adenocarcinomas including keratin 7, 8, and 18 (**Figure 2F, Figure S3D** and **Table S1**). Tumor cluster 2 (T2) showed significant enrichment of epithelial-to-mesenchymal transition markers including N-cadherin (*Cdh2*) and type I collagen genes (**Figure 2G, Figure S3D** and **Table S1**). Whether these two clusters represent intratumor heterogeneity of a single tumor clone or distinct intermixed independent tumors is not clear. These data suggest that RIT1^M90I^ expression in alveolar epithelial cells cooperates with loss of *Nf2* and *p53* to drive transdifferentiation to an EMT-like lung cancer state.

### RIT1 and YAP synergize to promote cJUN expression and EMT transcriptional programs

The Nf2 tumor suppressor is a critical upstream Hippo signal transducer and negative regulator of the oncoprotein YAP (Harvey et al. 2013). To model the synergy observed between RIT1 and Nf2 loss, we used human small airway lung epithelial cells (SALE) with ectopic expression of RIT1^M90I^ and activated YAP. SALE cells are immortalized, non-transformed human lung epithelial cells that we previously showed can be transformed by RIT1^M90I^ together with either *NF2* knockout or ectopic expression of YAP^8SA^, an engineered mutant of YAP with enhanced nuclear localization and activity (Vichas et al. 2021). Here we confirmed that combined RIT1^M90I^ and YAP^8SA^ expression synergized to promote growth in low attachment (Izar and Rotem 2016) compared to individual expression of each oncogene or the parental control cells (**Figure 3A**). To investigate the transcriptional programs underlying this phenotype, we analyzed bulk RNA-sequencing data from an isogenic series of SALE cells expressing RIT1^M90I^ and YAP^8SA^ either independently or in combination (**Figure 3B**) (Vichas et al. 2021). Both RIT1^M90I^ and YAP^8SA^ independently modulated transcriptional programs compared to parental cells, with 2185 and 46 differentially expressed genes (|log2FC|>2, p value < 0.01) compared to the vector control cells, respectively (**Table S2**). Furthermore, combined RIT1^M90I^/YAP^8SA^ expression markedly altered the transcriptome program in SALE, with induction and repression of hundreds of genes uniquely in the SALE-RIT1^M90I^/YAP^8SA^ (SALE-RY) co-expressed cells (**Figure 3B** and **Table S3**). To investigate the gene programs uniquely regulated under RIT1/YAP co-expression, we performed gene set enrichment analysis using the MSigDB database (Liberzon et al. 2015). Epithelial-to-mesenchymal transition (EMT) was the top hallmark pathway upregulated in SALE-RY cells compared to either vector control or cells expressing only activated YAP^8SA^ (**Figure S4A**). In pathway analysis of REACTOME pathways, alteration of extracellular matrix proteins was evident, with multiple cell adhesion and matrix pathways strongly upregulated in the SALE-RY cells compared to single-expressing or vector control cells (**Figure S4B**). Similar to the murine tumor cells, robust overexpression of multiple collagen genes was observed (**Figure S4C**).

**Figure 3.**
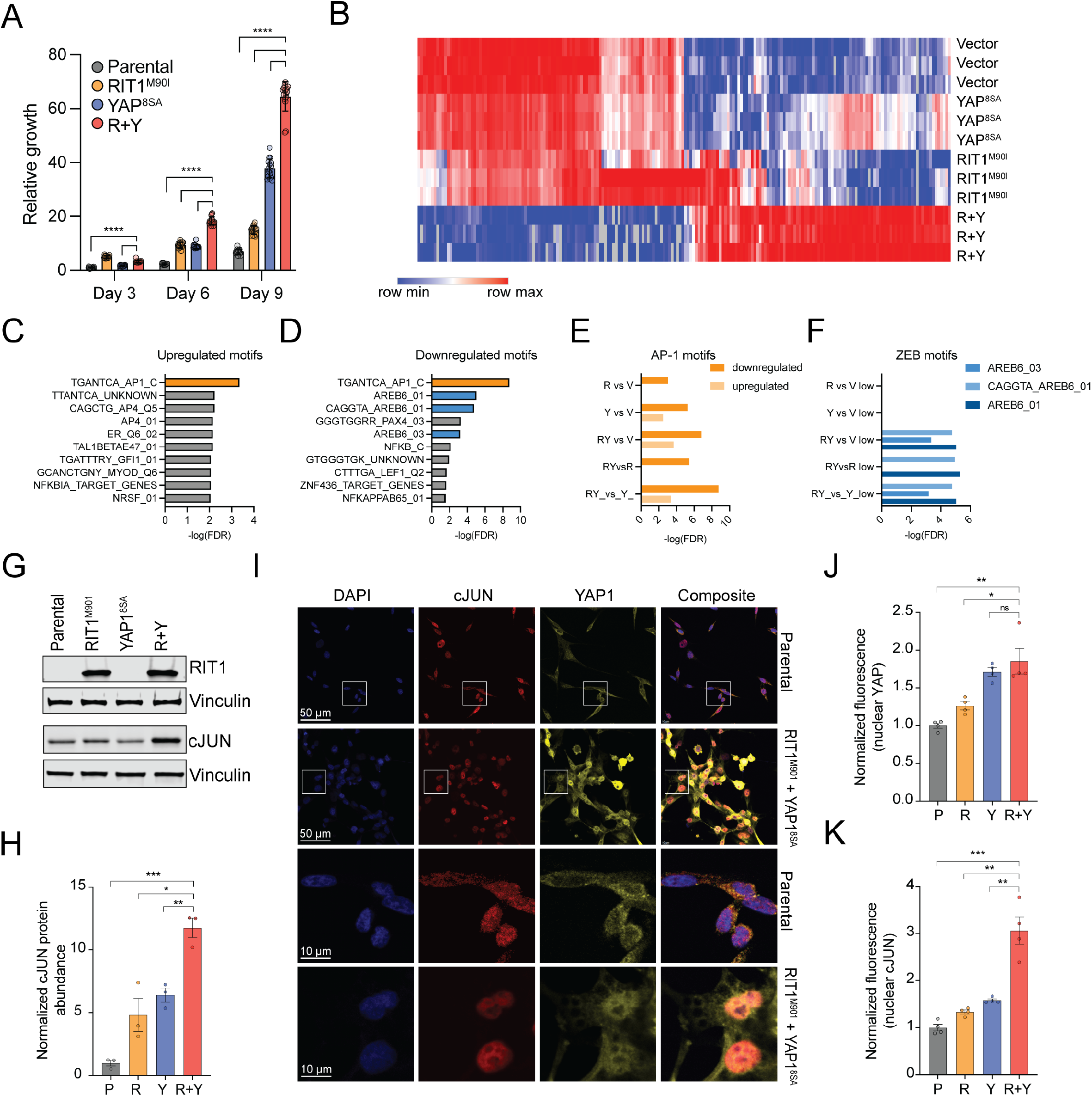
RIT1 and YAP synergize to promote cJUN expression and EMT. **A**, GILA assay of isogenic SALE cells expressing RIT1^M90I^ and YAP^8SA^ alone or in combination. 400 cells/well were seeding in a 96 well ultra-low attachment plate and growth was analyzed by CellTiterGlo after the days indicated. **B**, Heat map of gene expression data derived from bulk RNA-sequencing of isogenic SALE cells. Each column is a differentially expressed gene and each row is a replicate of the cell line variant indicated. The top 100 up- and down-regulated genes distinguishing R+Y from YAP^8SA^ cells were determined by marker selection based on the mean difference of R+Y replicates compared to YAP^8SA^ replicates and then genes and samples and genes were clustered using one minus Pearson correlation. See also **Table S2. C**, Motif enrichment of upregulated genes in R+Y compared to YAP^8SA^ cells using the MSigDB Transcription Factor Target (TFT) geneset. The orange bar indicates an AP-1 binding motif. **D**, Motif enrichment of downregulated genes in RY compared to Y cells using MSigDB TFT gene set. The orange bar indicates an AP-1 binding motif and the blue bars indicate AREB6/ZEB1 binding motifs. **E**, Comparison of GSEA significance across different comparisons of SALE isogenic cells for either upregulation or downregulation of genes containing the AP-1 binding motif TGANTCA. **F**, Comparison of GSEA significance across different comparisons of SALE isogenic cells for downregulation of genes containing any of the three AREB6/ZEB1 binding motifs shown. **G**, Western blot of lysates from isogenic SALE cells using antibodies against RIT1 and cJUN. Vinculin is used as a loading control. **H**, Quantification of biological replicates (n=3) of cJUN abundance determined by western blot, normalized to loading control and parental cJUN abundance. **I**, Immunofluorescence and confocal images of SALE cells plated onto collagen-coated glass slides, and fixed and stained with anti-YAP and anti-cJUN antibodies and counterstained with DAPI. In the top panels, white boxes indicate the regions of interest shown at higher magnification in the bottom two panels. **J**, Quantification of nuclear YAP fluorescence intensity determined by confocal microscopy from biological replicates (n=3-4) of isogenic SALE cells. **K**, Quantification of nuclear cJUN fluorescence intensity determined by confocal microscopy from biological replicates (n=3-4) of isogenic SALE cells. ns, not significant; *, p < 0.05, **, p < 0.01, ***, p < 0.001, ****, p < 0.0001 by unpaired two-tailed t-test.

The extensive rewiring of the transcriptional program in SALE-RIT1^M90I^/YAP^8SA^ cells suggested to us that RIT1^M90I^ might alter transcriptional regulation. To identify transcription factors involved in the oncogenic phenotype of SALE-RY cells, we performed motif enrichment analysis of differentially expressed genes in SALE-RY cells. Compared to SALE cells expressing YAP^8SA^ only, genes both up- and down-regulated in the SALE-RY cells were significantly enriched for the TGANTCA motif of the AP-1 transcription factor cJUN (**Figure 3C-D**). Both RIT1^M90I^ and YAP^8SA^ alone to some extent modulated AP-1 target genes, but SALE-RY cells has the strongest enrichment of AP-1 sites in downregulated genes among the cell comparisons (**Figure 3E**) suggesting altered AP-1 activity in the setting of combined RIT1^M90I^/YAP^8SA^ activation. Suppression of genes containing the EMT master regulator AREB6 (ZEB1) binding sites were significantly enriched in SALE-RY cells relative to vector control (**Figure 3F**) and further enriched relative to cells with single expression of either RIT1^M90I^ or YAP^8SA^ (**Figure 3F**). ZEB1 is a transcriptional repressor involved in initiation of EMT and suppression of epithelial lineage factors such as E-cadherin. Western blot of SALE isogenic cells showed marked phosphorylation of ERK and AKT in SALE-RY cells (**Figure S4D**), as well as significantly increased protein abundance of cJUN itself (**Figure 3G-H**). Immunofluorescence of YAP and cJUN confirmed increased nuclear YAP staining in the YAP^8SA^ cells (**Figure 3I-J**) and increased cJUN nuclear expression in SALE-RY cells compared to cells individually expressing RIT1^M90I^ or YAP^8SA^ (**Figure 3I, 3K**). Together these data suggest that RIT1^M90I^ and YAP^8SA^ cooperate to promote EMT transcriptional programs through modulation of cJUN protein abundance and transcriptional activity.

### Combined MEK and TEAD inhibition suppresses growth of RIT1^M90I^-mutant lung cancer cells

RIT1 is a noncanonical RAS GTPase which shares with canonical RAS proteins the ability to activate MEK/ERK signaling via BRAF (Van et al. 2020) or CRAF (Castel et al. 2019). cJUN abundance is responsive to upstream mitogenic growth factor signaling by ERK and JNK and has been linked to drug resistance and phenotype switching in *BRAF*-mutant melanoma (Ramsdale et al. 2015; Karin 1995). We therefore tested whether MEK inhibition with the small molecule trametinib (Gilmartin et al. 2011) could suppress the oncogenic growth of SALE-RY cells, alone or in combination with VT104 (Tang et al. 2021), an analog of VT3989, a pan-TEAD autopalmitoylation inhibitor that suppresses TEAD-mediated transcription and has shown encouraging results in clinical development (Yap et al. 2023). Increasing doses of trametinib and VT104 treatment completely prevented growth in low attachment of SALE-RY cells, confirming that MEK and TEAD are important mediators of the oncogenic synergy of RIT1^M90I^ and YAP^8SA^ (**Figure 4A**). S-RPN-1 cells, derived from a murine Ad5mSPC-Cre/*RIT1*^M90I^/*p53*^*fl/fl*^*/Nf2*^*fl/fl*^ lung tumor were sensitive to single agent trametinib treatment (IC50 = 0.426 μM), single agent TEAD inhibition with VT104 (IC50 = 0.397 μM) or VT107 (IC50 = 0.227 μM), and to a lesser extent, TEAD1 inhibition with VT103 (IC50 = 4.66 μM). Importantly, a less active enantiomer of VT104, VT106, showed markedly reduced activity (IC50 > 9 μM) (**FIgure 4B**). Combined trametinib and TEAD inhibition of either pan-TEAD (VT104 or VT107) or TEAD1 (VT103) provided the best cell growth inhibition (**Figure 4C**, range 43-57 nM). These data demonstrate cooperativity between MEK and TEAD inhibition in S-RPN-1 cells and suggest that combined inhibition of TEAD and MEK might provide a promising avenue for treatment of *RIT1*-mutant lung cancer.

**Figure 4.**
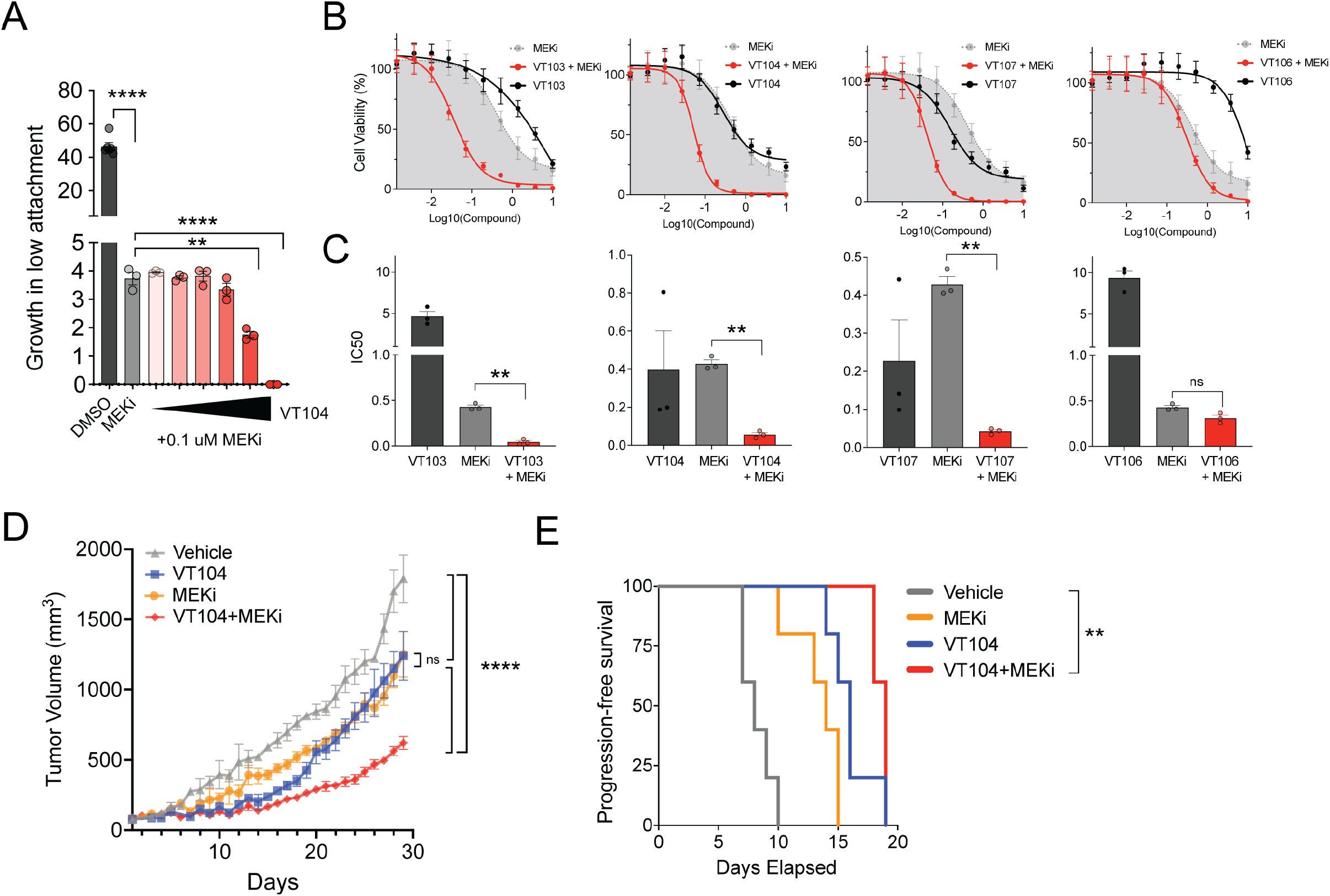
Combined MEK and TEAD inhibition suppresses growth of RIT1^M90I^-mutant lung cancer cells. **A**, Growth-in-low attachment assay of SALE-RY cells in the presence and absence of 0.1 μM trametinib (MEKi) or increasing dosages of the pan-TEAD autopalmitoylation inhibitor, VT104 (0.1, 0.25, 0.6, 1.6, 4, and 10 μM). Cells were plated on ultra-low attachment plates and incubated with each drug for 7 days before cell growth was determined by CellTiterGlo. The data shown is the mean and s.e.m. of three technical replicates. **B**, In vitro dose response assay of S-RPN-1 tumor cells with trametinib (MEKi) or the TEAD1-selective inhibitor VT103, pan-TEAD inhibitors VT104 and VT107, and less-active pan-TEAD inhibitor enantiomer VT106. Cells were plated in 96 well plates and analyzed by CellTiterGlo after 7 days of drug treatment. The data shown is the mean and s.e.m. of three independent biological replicates. **C**, IC50 of compounds tested in (B). The data shown is the mean and s.e.m. of the IC50 determined from three independent experiments. **D**, Allograft tumor growth of S-RPN-1 cells in the flanks of immunocompromised mice (n=5 per group). Mice were enrolled in the study when tumor reached >70 mm^3^ after which animals were randomized to the treatment groups indicated. See Methods for dosing details. **E**, Kaplan-Meier survival analysis of progress-free survival (PFS) of the mice shown in (D). PFS was defined as a 300% tumor volume increase. ns, not significant; *, p < 0.05, **, p < 0.01, ***, p < 0.001, ****, p < 0.0001 by unpaired two-tailed t-test (A), paired two-tailed t-test (C), one-way ANOVA with multiple testing correction (D), or log-rank test (E).

To test this hypothesis in vivo, we implanted allografts of S-RPN-1 cells into the flanks of immunocompromised host animals and treated mice for ∼1 month with either single agent VT104, trametinib, or combination VT104/trametinib. Both trametinib and VT104 had similar single agent activity, each providing 30% tumor growth inhibition at day 29 (adjusted p < 0.0001 by one-way ANOVA). Combination treatment of VT104 and trametinib substantially reduced tumor growth, providing 65% tumor growth inhibition (**Figure 4D;** adjusted p < 0.0001 compared to either single agent by one-way ANOVA) and extension of progression-free survival (**Figure 4E**). These data confirm the critical role of both MEK and TEAD in the growth of *RIT1*^M90I^/*p53*^*fl/fl*^*/Nf2*^*fl/fl*^ tumor cells and demonstrate the promise of combined MEK/TEAD inhibition for treatment of *RIT1-*mutant lung cancer.

## DISCUSSION

RIT1 shares ∼50% amino acid identity with canonical Ras proteins HRAS, KRAS, and NRAS (Reuther and Der 2000; Lo et al. 2021). Germline variants in *RIT1* underlie the RASopathy, Noonan syndrome, also caused by germline variants in *KRAS* and *NRAS*. However, unlike canonical Ras genes which are among the most frequently mutated genes in cancer, *RIT1* is mutated only in 1-2% of various solid cancers and myeloid malignancies. The biological mechanisms that may explain this difference are not understood, and insight into the mechanisms governing oncogenic activity of Ras family GTPases have important implications for tumor therapy, as inhibition of Ras GTPases for cancer therapy has regained attention with the recent successful development of direct inhibitors of KRAS^G12C^ (Hong et al. 2020; Jänne et al. 2022; Awad et al. 2021).

*RIT1* mutations in lung cancer cluster near the switch II domain which is believed to be involved in protein-protein interactions and effector binding (Berger et al. 2014). *RIT1* mutations in lung cancer are mutually exclusive with *KRAS* mutations and, like *KRAS* mutations, can confer resistance to EGFR inhibition and promote cell transformation (Berger et al. 2016, 2014). Due to the extensive similarity in structure and function between RIT1 and Ras proteins, it is therefore surprising that RIT1^M90I^ expression alone was insufficient to promote a significant tumor growth phenotype and showed little cooperation with *p53* loss. One potential explanation for this disconnect between Ras and RIT1 activity in vivo could be the importance of RIT1 protein abundance to its function (Castel et al. 2019) and the potentially weak expression of transgenic RIT1^M90I^ driven by the endogenous Rosa26 promoter. RIT1 protein level is believed to be key to its oncogenic function since the M90I variant disrupts LZTR1-mediated degradation of RIT1 (Castel et al. 2019), and wild-type RIT1 overexpression phenocopies mutant RIT1 (Lo et al. 2021). Thus modeling RIT1^M90I^ expression from its endogenous promoter or via transgenic overexpression might lead to increased protein expression and accelerated RIT1^M90I^-driven tumorigenesis.

Another explanation for the mild lung cancer phenotype driven by RIT1^M90I^ alone could be the requirement for cooperating genetic factors. This explanation is also consistent with the rare mutational frequency of *RIT1* in cancer. If *RIT1* mutations only confer a selective advantage to lung cells in the context of other genetic events, then the frequency of *RIT1* mutations observed in cancer would be lower than those mutations that are sufficient to promote a selective advantage on their own such as *KRAS*. We previously identified Hippo pathway inactivation and YAP nuclear accumulation as common features of human *RIT1*-mutant lung adenocarcinoma (Vichas et al. 2021). Therefore in the present work, we tested whether modeling combined *RIT1*^M90I^/*p53*^*fl/fl*^*/Nf2*^*fl/fl*^ mutation would result in enhanced tumor growth in vivo. Incredibly, this combined mutation resulted in lethal lung tumor development in only weeks, confirming the unique oncogenic synergy between RIT1^M90I^, p53 loss, and Hippo inactivation.

The Hippo signaling pathway is a tumor suppressive pathway that links signals from the external cell adhesion matrix to cell growth decisions mediated by downstream YAP/TEAD activation. Because RIT1, like Nf2, is known to bind to the plasma membrane (Van et al. 2020), it seems possible that RIT1 might impart its regulation of Hippo/YAP activity near the membrane or in the cytosol. However, it was surprising that mutant RIT1 cooperated not only with upstream Hippo inactivation via *Nf2* loss, but also with the constitutively active YAP^8SA^ variant. YAP^8SA^ lacks all of the phosphorylation sites used by LATS kinases to phosphorylate YAP and retain it in the cytosol for degradation and should therefore be insensitive to upstream changes in Hippo pathway regulation. This knowledge led us to hypothesize that the mechanism of synergy between RIT1 and YAP lies somewhere in the nucleus, likely through transcriptional regulation. In support of this hypothesis, RNA-seq of both the murine tumor model and SALE-RY cells revealed altered regulation of genes with binding sites for the AP-1 transcription factor cJUN, which we showed was overexpressed in SALE-RY cells. These data link RIT1 to an AP-1 transcriptional program shown to alter chromatin binding of the TEAD transcription factors which transmit YAP’s oncogenic signaling (Zanconato et al. 2015).

The tumors in *RIT1*^M90I^/*p53*^*fl/fl*^*/Nf2*^*fl/fl*^ mice were highly undifferentiated or mesenchymal in phenotype. Unexpectedly, this mesenchymal-like tumor could be driven by RIT1^M90I^ activation in AT2 cells, the believed cell of origin of lung adenocarcinoma (Ferone et al. 2020). We propose a model that can reconcile this paradox in combined *RIT1*^M90I^/*p53*^*fl/fl*^*/Nf2*^*fl/fl*^ mutation in AT2 cells promotes both tumor formation and a cell lineage transition resembling EMT. This model is in agreement with a growing understanding that lung squamous cell carcinoma, lung adenocarcinoma, and small cell lung cancer can all be created from different cells of origin in murine models (Ferone et al. 2020). Moreover, in both human prostate cancer and lung cancer, it is now well established that tumors can develop resistance to androgen receptor or EGFR-targeted therapies via histologic transformation to neuroendocrine cancer (Rubin et al. 2020). Thus, we now understand that apparent tumor cell state is not linked with fidelity to cell-of-origin.

Despite decades of concerted effort in establishment of cancer cell line models from the Cancer Cell Line Factory (Boehm and Golub 2015), Cancer Dependency Map (Tsherniak et al. 2017), and others (Gazdar et al. 2010), representation of rare molecular subtypes of cancer such as *RIT1*-mutant lung cancer is limited. Yet cell line and patient-derived xenograft models are essential for discovery and pre-clinical validation of new strategies for cancer therapy. Our development of a new autochthonous model of RIT1^M90I^-mutant lung cancer enabled us to identify Hippo pathway inactivation as a critical synergistic event with oncogenic RIT1^M90I^. These functional data, combined with the frequent YAP activation and Hippo inactivation observed in human *RIT1*-mutant lung cancer (Vichas et al. 2021), nominate MEK and TEAD targeting as a new approach for cancer therapy of *RIT1*-mutant lung cancer.

## METHODS

### Generation of the LSL-RIT1^M90I^ mouse model

LSL-RIT1^M90I^ genetically engineered mice were generated by homologous recombination of a targeting cassette containing 5’ and 3’ homology arms to the mouse Rosa26 locus and carrying an insert consisting of human RIT1 cDNA with a ATG>ATA p.M90I mutation. The targeting cassette was generated by cloning the human RIT1 p.M90I cDNA sequence first into the pBTG plasmid using NheI and XhoI sites. pBTG (pBigT-IRES-GFP; DM#268) was a gift from Douglas Melton (Addgene plasmid # 15037; http://n2t.net/addgene:15037 ; RRID:Addgene_15037). The RIT1-IRES-GFP cassette was then subcloned into the pRosa26PAm1 backbone by restriction cloning using PacI and AscI sites. pRosa26PAm1 (DM#272) was a gift from Douglas Melton (Addgene plasmid #15036;http://n2t.net/addgene:15036 ; RRID:Addgene_15036). The targeting vector was verified by restriction digest and sequencing. Gene targeted ES cells and chimeric mice were created at the Brigham and Women’s Hospital Transgenic Core Facility. The construct was linearized using MluI and electroporated into CJ7 cells, followed by positive selection with neomycin and negative selection with diphtheria toxin. Selected clones were screened by three PCR assays to ensure proper recombination. First, presence/absence of the neomycin cassette was assessed with primers specific for the neomycin cassette. Second, correct integration was confirmed using PCR from the insert across the 5’ arm and into the endogenous locus. Last, correct integration on the 3’ end was assessed using PCR from the insert across the 3’ arm into the 3’ endogenous locus. Clones 5 and 6 were selected for chimera generation. Chimeras from both clones were crossed to wild-type C57BL/6 mice and both clones had successful germline transmission of the LSL-RIT1^M90I^ allele. Offspring were genotyped by analysis of tail or ear punch tissue using genotyping assays at Transnetyx. Animals generated from clone 5 ES cells were used for all subsequent experiments and were maintained on a mixed 129/B6 genetic background. To facilitate transfer from the BWH Transgenic Core to the Broad Institute, additional mice were generated by in vitro fertilization at Charles River Laboratories. Breeding and experimental animals were then transferred to Fred Hutchinson Cancer Center, with the experiments described performed at Fred Hutchinson Cancer Center.

### Verification of Cre delivery using mTmG mice

6-week-old mTmG females (JAX strain 007676) were obtained from Jackson Laboratory, Bar Harbor; ME, USA, and allowed to acclimate to the facility for at least 1 week. mTmG mice have a tdTomato reporter (mT) that is expressed in all tissue types and upon delivery of Cre is converted to EGFP (mG). About 9 weeks after the initial administration, lungs from the mTmG mouse model were harvested and the postcaval lobe and the left lung were fixed in 10% formalin, transferred to a 30% sucrose solution, embedded in OCT, and then sent to the histology core for analysis. Cells were extracted from the superior, middle, and inferior lobes and were isolated for flow cytometry. DAPI was added to the cells for flow cytometry and each sample from the mice was analyzed in FloJo.

### Mouse experiments and strains

All animal experiments were carried out with approval by and in accordance with the ethical guidelines of the Fred Hutchinson Cancer Research Center Institutional Animal Care and Use Committee (Protocol #50967, PI: Berger). Experiments were performed in the Fred Hutch Comparative Medicine facility, which is fully accredited by the Association for Assessment and Accreditation of Laboratory Animal Care (AAALAC) and complies with all United States Department of Agriculture (USDA), Public Health Service (PHS), Washington State and local area animal welfare regulations. For analysis of mice with germline activation of RIT1^M90I^, LSL-RIT1^M90I^ mice were crossed with hemizygous mice expressing Cre under the SRY-box containing gene 2 promoter (Hayashi et al. 2002) (Jackson Laboratory #008454). This line reportedly harbors some Cre recombinase activity in Cre-zygotes when delivered from female Cre+ breeders (Hayashi et al. 2003), although consistent recombination Cre-offspring from Cre+ dams was not observed. For analysis of lung tumor initiation, *Rosa26-LSL-RIT1*^*M90I*^ were crossed with mice containing a conditional *Trp53* allele (Marino et al. 2000) (Jackson Laboratory #008462) or a conditional *Nf2* allele (Riken strain RBRC02344) (Giovannini et al. 2000).

### Intratracheal adenovirus delivery

10-12 week old mice were anesthetized with Avertin and virus delivered as previously described (DuPage et al. 2009). Ad5CMV-Cre and Ad5mSPC-Cre virus was obtained from the University of Iowa Viral vector core and administered intratracheally in 75μL of a DMEM and CaCl_2_ solution containing 1.13*10^10^ PFU/mL adenovirus.

### MicroCT imaging

Lung CT scans were acquired via respiratory-gated high-speed micro-CT lung 3D data sets (Quantum GX2 micro-CT, Perkin Elmer) with 90 kV tube voltage, 88 μA tube current, 36mm FOV, 72 μm voxel size, and a total scan time of 4 minutes per mouse. All data analyzed was from the expiratory dataset resulting from the respiratory-gated output. A Tungsten anode is the X-ray source in this scanner and a fixed filter of 0.5 mm Aluminum (Al) and 0.06 mm Copper (Cu) is placed in front of the exit port to remove low-energy X-rays that contribute to dose but do not improve image quality. One scan gives a radiation dose of 912 mGy.

### Histopathology and immunohistochemistry

Lungs were harvested after PBS perfusion and inflation with low melting point agarose. Lungs were fixed in 10% formalin for 24-48 hours while rocking at 4 ºC before embedding in paraffin blocks and sectioning at 4-5 μm onto charged slides. Slides with paraffin sections were baked at 60°C, processed using an automated Stainer (Sakura Tissue-Tek Prisma) for Hematoxylin and Eosin (H&E) staining, and coverslipped with permanent mounting media. Whole slides stained with H&E were scanned using TissueFAXs slide scanner at 2.5x/0.075 (air) and then the field of view (FOV) was taken at 10x/0.3 (air) then stitched together (TissueGnostics). IHC was performed on a Leica Bond RX automated stainer (Leica Biosystems, Buffalo Grove, IL) using Leica Bond reagents. Endogenous peroxidase was blocked with 3% H2O2 for 5 min followed by protein blocking with TCT buffer (0.05M Tris, 0.15M NaCl, 0.25% Casein, 0.1% Tween 20, and 0.05% Proclin 300 at pH 7.6 +/-0.1) for 10 min. Heat induced epitope retrieval was performed in a pH 9.0 buffer. Primary antibodies TTF1 (Abcam #ab75013, 1:250) or Ki67 (Cell Signaling #12202, 1:2000) were incubated for one hour and secondary polymers (Rabbit Polymer HRP) were applied, followed by Mixed Refine DAB (Leica DS9800) for 10 min and counterstained with Refine Hematoxylin (Leica DS9800) for 4 min after which slides were dehydrated, cleared and coverslipped with permanent mounting media. Pathology and tumor incidence was determined by two board-certified veterinary pathologists (AK and RG). Each tissue was examined for lesion category including pulmonary hyperplasia, adenoma, adenocarcinoma, and undifferentiated carcinoma, and received a score of 0-2 for each lesion category (Score 0: no tumor or pulmonary hyperplasia lesion, Score 1: solitary tumor or pulmonary hyperplasia lesion, Score 2: multiple tumors or pulmonary hyperplasia lesions).

### Single-cell RNA-sequencing of RIT1^M90I^-mutant lung cancer

A *RIT1*^M90I^/*p53*^*fl/fl*^*/Nf2*^*fl/fl*^ mouse was euthanized 20 weeks after Ad5mSPC-Cre treatment. Following euthanasia, the mouse was perfused with PBS and the lungs were removed and a visible tumor resected. This isolated tumor was dissociated using the Tumor Dissociation Kit, mouse (Miltenyi Biotech, catalog #130-096-730) and gentleMACS “C” columns (Miltenyi Biotech, catalog #130-093-237) following the suggested manufacturer’s protocol. The dissociated cells were pelleted, resuspended in RPMI 1640 (Gibco, catalog #11875119) and filtered through the MACS SmartStrainer (Miltenyi Biotech, catalog #130-098-462). The resulting cell suspension was centrifuged at 300 rcf for 7 minutes at room temperature, followed by supernatant removal and resuspension of cell pellet in Red Blood Lysis Solution (Fisher Scientific, catalog #AAJ62150AP). Cells were mixed and incubated at 4 ºC for 3 min. Cells were then washed with Wash Buffer (0.04% BSA (ThermoFisher, catalogue #B9000S) in 1x DPBS (Corning, catalogue #21-030-CVR)) and centrifuged again at 300 rcf for 4 min. Supernatant was removed and the cells were resuspended again in appropriate amount of Wash Buffer 0.04% BSA (ThermoFisher, catalogue #B9000S) in 1x DPBS (Corning, catalogue #21-030-CVR) followed by cell counting for use in single cell library preparation. Single cell sequencing libraries were generated using the Chromium Next GEM Single Cell 3′ Library and Gel Bead kit v3.1 (Dual Index) as per the manufacturer’s protocol. Two libraries were prepared from the same tumor with targeted recovery of 10,000 cells/reaction. Libraries were sequenced on a NextSeq 2000 (Illumina) at the Fred Hutch Genomics Shared Resource. The reads generated after sequencing were aligned in CellRanger v7.1.0 to a modified mm10 mouse reference in which we added the RIT1^M90I^ transgene reference sequence. All further analysis of the resulting count matrices was performed in Python (v3.9) following the standard scanpy pipelines. Cells were excluded if they had below 100 genes. Similarly genes were excluded if present in fewer than 3 cells. Further filtering for doublets was done using scVI. Cells having >20% of reads as mitochondrial reads were also excluded. After filtering, 11,241 cells were obtained for analysis from two 10X Chromium reactions. Count data was normalized with the total target sum of 10,000 per cells and log transformation. This was followed by selecting 3000 highly variable genes and then by integration of the two libraries using scVI. Finally, a neighborhood graph was constructed with cosine as the similarity matrix and the dataset was clustered using Leiden clustering and UMAP visualization. The final clustering led to formation of 13 separate subsets which were manually verified for proper separation for different cell types. For each cluster, the top 100 ranked genes compared to other clusters were identified using the Wilcoxon test method. PanglaoDB was queried using these genes to find the top predicted cell type, and some of these top ranked genes were labeled as marker genes after referring to the literature. Some of the subsets belonged to the same cell type, which reduced the number of different clusters/cell types to 11. The following are the marker genes used to identify the cell types: AT & Club cells: Ager, Sftpc, Scgb1a1; Alveolar Macrophages: Ear1 and Ear2; B-cells: Cd79b, Ms4a1; Endothelial cells: Pecam1, Egfl7; Fibroblasts: Col3a1, Dcn; Macrophages: Ccl9, F13a1; Neutrophils: S100a8, S100a9; T-cells: Cd3g, Cd3e. The remaining two clusters we named “T1” and “T2” were identified as the tumor cell clusters with high expression of transgenic RIT1.

### SALE RNA-sequencing analysis and motif enrichment

Bulk RNA-sequencing of isogenic SALE cells (parental, RIT1^M90I^, YAP^8SA^, and RIT^M90I^+YAP^8SA^) have been previously described and are available from the NCBI Gene Expression Omnibus (GEO) under dataset ID #GSE165631. Note that in prior work and at Addgene, the YAP^8SA^ mutant has also been referred to as YAP^5SA^. Differentially expressed genes in each comparison were determined using the Broad Institute’s Morpheus software (https://software.broadinstitute.org/morpheus/) marker enrichment tool by the mean difference method. The top 100 upregulated or top 100 downregulated genes were used to query MSigDB using the Investigate Gene Sets Gene Set Enrichment Analysis (GSEA) method after excluding the ectopically expressed RIT1 and YAP1 genes.

### Antibodies

Primary antibodies used for immunoblotting: β-Actin 1:1000 (Cell Signaling Technology, 4970), Vinculin 1:10000 (Sigma, V9264), RIT1 1:2000 (Abcam, Ab53720), p44/42 MAPK 1:1000 (Erk1/2) (Cell Signaling Technology, 9107), Phospho-p44/42 MAPK (Erk1/2) 1:1000 (Thr202/Tyr204) (Cell Signaling Technology, 4370), AKT 1:1000 (Cell Signaling Technology, 2920), Phospho-AKT 1:1000 (Cell Signaling Technology, 9112), YAP 1:200 (Cell Signaling Technology, D8H1X), cJUN 1:1000 (Cell Signaling Technology, 9165). Primary antibodies for immunohistochemistry: Desmin (Dako M0760 D33 1:50), Vimentin (Novus NB300-223 1:450), SMA (Dako M0851 A23 1:150), Mesothelein (MyBioSource MBS2528505 H50 1:1600), Pan-cytokeratin (Abcam ab9377 1:500).

### Immunofluorescence

Cells were plated on collagen coated coverslips in a 6 well plate. After 24 hours, the cells were rinsed with PBS, fixed with 4% paraformaldehyde, permeabilized with 0.25% Triton X-100 in PBS, and then blocked in 1% BSA in PBS + 0.1% Tween20 (A9647-10G Millipore) for one hour. Primary antibodies were diluted in 50 ul 1% BSA PBST with 1:200 cjUN (60A8) (Cell Signaling Technology, 9165) and 1:50 YAP (Santa Cruz Biotechnology, sc-101199) and incubated at 4°C overnight. Cover slips were washed in PBST and then incubated in secondary antibody for an hour. Cells were then mounted with ProLong™ Diamond Antifade Mountant with DAPI (Fisher Scientific) and left overnight before imaging using a Stellaris8 confocal microscope (Leica).

### In vitro dose response assays

S-RPN-1 or SALE cells were plated in 96 well plates using the Multidrop Combi Reagent Dispenser (Thermo Fisher). After 24 hrs, cells were treated with TEAD or MEK inhibitors in a serial dilution using the D300e dispenser (Tecan). 7 days after treatment, 40 ul of CellTiterGlo reagent (Promega) was added and cell viability was quantified on the LUMIstar Omega microplate reader (BMG Labtech). The fraction of cell viability was calculated from normalizing the treated cells to the DMSO control. Graphs were constructed using GraphPad Prism v10.2.3 non-linear curve fit to assess viability. Trametinib was obtained from SelleckChem (S2673) and VT compounds were provided by Vivace therapeutics: VT103, VT104, VT106, VT107.

### Pre-clinical study in S-RPN-1 allografts

To construct the allografts, 0.2 million S-RPN-1 cells were implanted subcutaneously into 1 flank of 20 NU/J mice. When the tumor volume reached 70 mm^3 mice were enrolled in the experiment. VT104 was provided by Vivace Therapeutics and was formulated daily in 5% DMSO, 10% Solutol, and 85% D5W. Trametinib was purchased from Selleckchem and formulated weekly in 4% DMSO + corn oil. Five animals per arm were dosed on a 5 day on/2 day off schedule via daily oral gavage with vehicle control, 10 mg/kg/day VT104, 0.5 mg/kg/day trametinib, or combination of VT104/trametinib dosed 30 minutes apart. Statistical comparison of the four arms was performed using a One-way ANOVA comparing all conditions to all other conditions using the mean of each arm on each day with multiple hypothesis correction using the Tukey method with alpha of 0.05.

## ACKNOWLEDGEMENTS

We thank Dr. Matthew Meyerson for providing laboratory space and support for the initial generation of RIT1 mice. We thank the Brigham and Women’s Hospital Transgenic core for technical services relating to generation of RIT1 mice. We thank Tracy Tang and Len Post from Vivace Therapeutics for advice and discussion and providing the VT compounds. This research was supported by the Genomics Shared Resource, Experimental Histopathology Shared Resource RRID:SCR_022612, Cellular Imaging Shared Resource, Comparative Medicine Shared Resource RRID:SCR_022610 and Preclinical Imaging Shared Resource of the Fred Hutch/University of Washington/Seattle Children’s Cancer Consortium (P30 CA015704). This work was funded in part by NIH/NCI grants R00CA197762 and R37CA252050 to AHB, NIH/NCI Cancer Center Support Grant P30CA015704, and NIH/NCI Lung SPORE grant P50CA228944.

## AUTHOR CONTRIBUTIONS

AHB conceived of the study, designed experiments, and obtained funding. MCR, SG, SM, MM, SK, SO, AW, FD, ARL, and EC designed and performed experiments and analyzed data. SG, SM, and AL performed computational analyses. RG and AK performed expert pathology review. AHB, MCR, SG, and SM generated the figures. AHB wrote the paper. All authors read, edited, and approved the paper.

