## Supplementary figures and images for "Mutant RIT1 cooperates with YAP to drive an EMT-like lung cancer state"

### Supplemental Figures

Rominger Figure S1

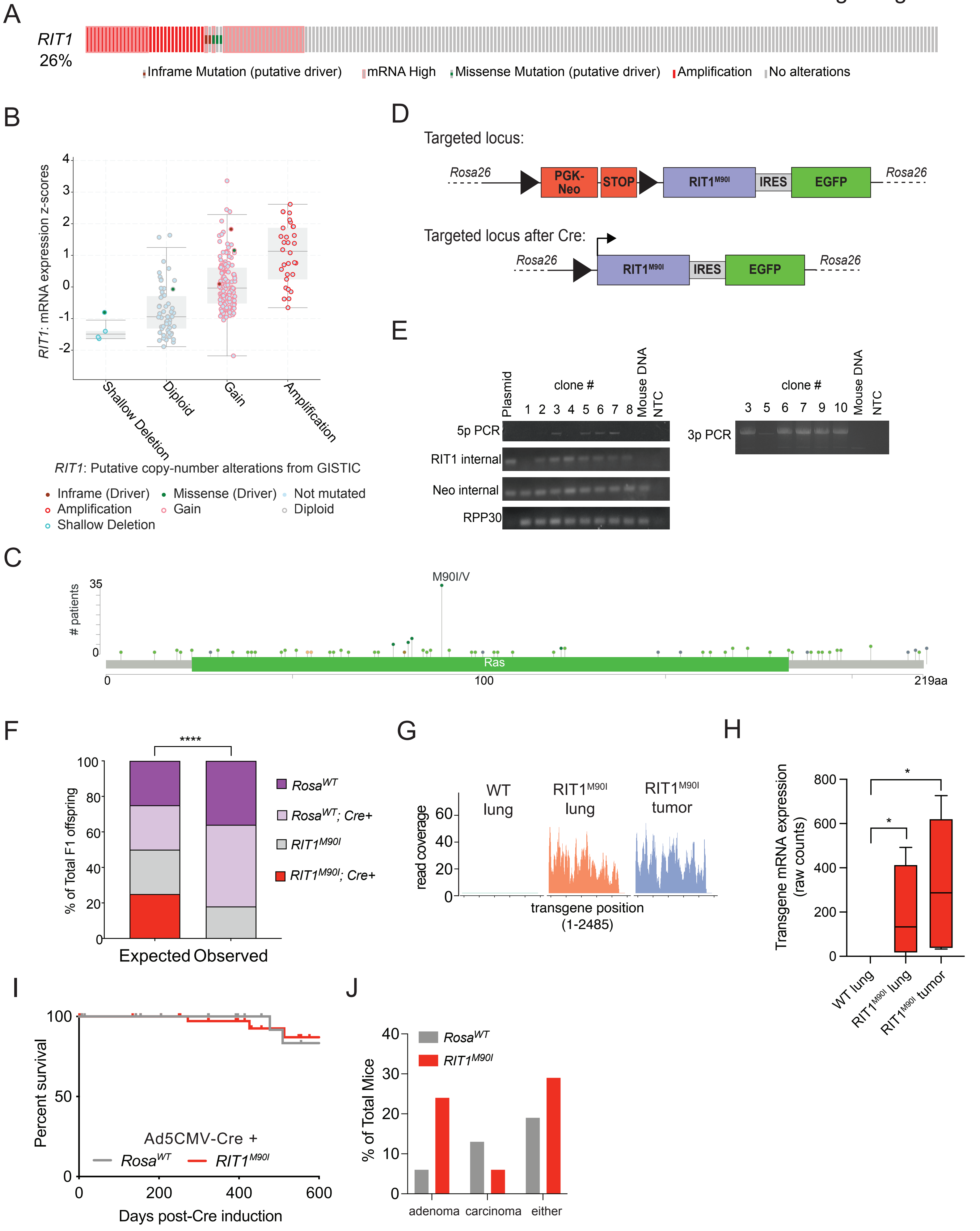

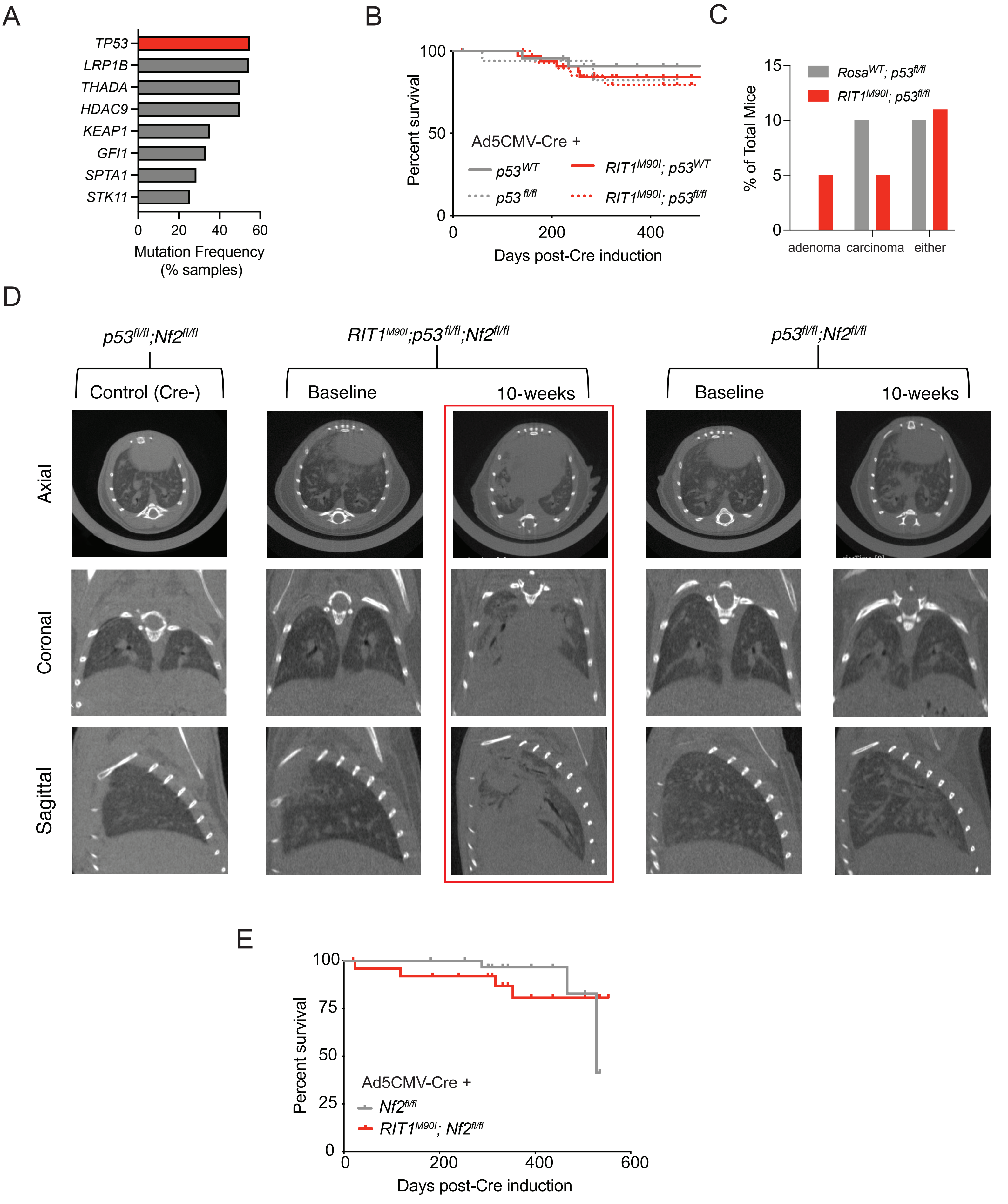

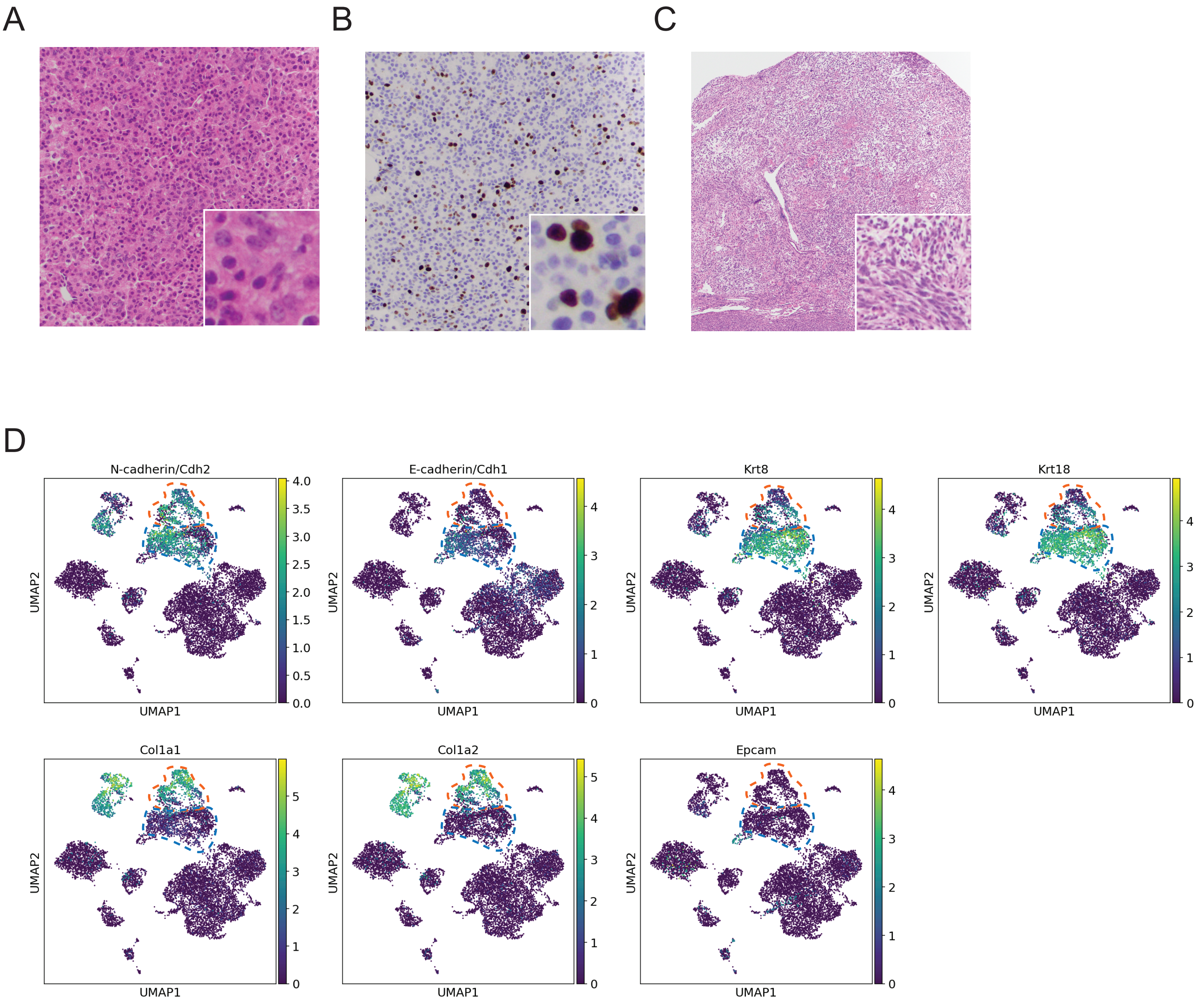

Rominger Figure S4

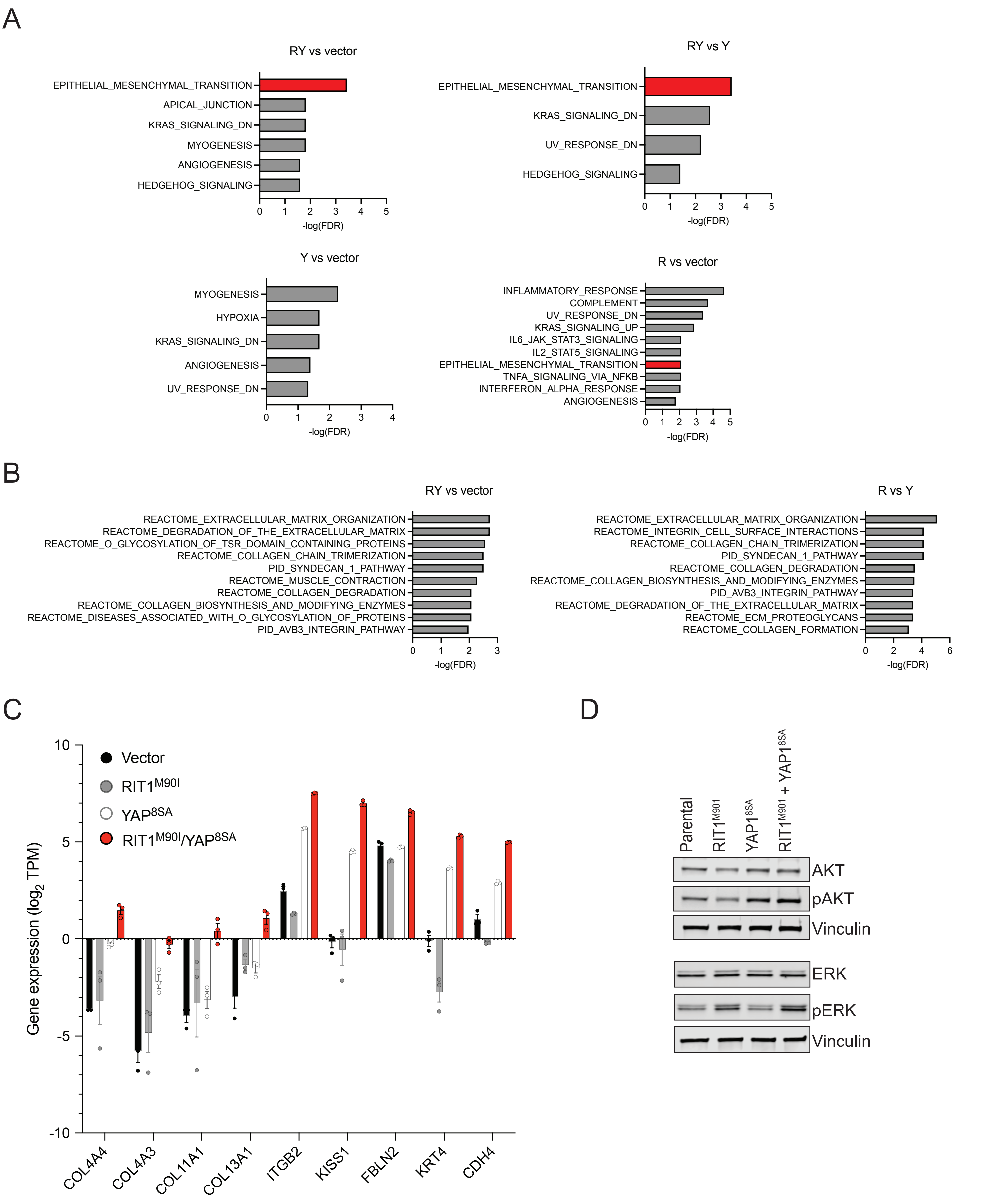
